# Impacts of agrochemical intensification on the assembly and reassembly of a mainland-island model metacommunity

**DOI:** 10.1101/2021.02.23.432475

**Authors:** Rodolfo Mei Pelinson, Bianca Rodrigues Strecht, Erika Mayumi Shimabukuro, Luis Cesar Schiesari

## Abstract

Many lentic aquatic environments are found embedded in agricultural fields, forming complex metacommunity structures. These habitats are vulnerable to contamination by agrochemicals, which can differentially affect local communities depending on the intensity and variability of species dispersal rates. We conducted a field experiment to assess how agrochemical intensification simulating the conversion of savannas into managed pastures and sugarcane fields affects freshwater community structure at different levels of spatial isolation. We constructed forty-five 1,200-L artificial ponds in a savanna landscape at three distances from a source wetland (30 m, 120 m, and 480 m). Ponds were spontaneously colonized by aquatic insects and amphibians and treated with no agrochemicals (‘savanna’ treatment), fertilizers (‘pasture’ treatment), or fertilizers and a single pulse of the insecticide fipronil and the herbicide 2,4-D (‘sugar cane’ treatment) following realistic dosages and application schedules. The experiment encompassed the entire rainy season. ‘Pasture’ communities were only slightly different from controls largely because two predatory insect taxa were more abundant in ‘pasture’ ponds. ‘Sugarcane’ communities strongly diverged from other treatments after the insecticide application, when a decrease in insect abundance indirectly benefitted amphibian populations. However, this effect had nearly disappeared by the end of the rainy season. The herbicide pulse had no effect on community structure. Spatial isolation changed community structure by increasing the abundance of non-predatory insects. However, it did not affect all predatory insects nor, surprisingly, amphibians. Therefore, spatial isolation did not change the effects of agrochemicals on community structure. Because agrochemical application frequently overlaps with the rainy season in many monocultures, it can strongly affect temporary pond communities. Ponds embedded in pastures might suffer mild consequences of fertilization by favoring the abundance of few predators through *bottom-up* effects. Ponds in sugarcane fields, however, might experience a decline in the insect population, followed by an increase in the abundance of amphibians tolerant to environmental degradation. Furthermore, we found no evidence that isolation by distance can change the general effects of chemical intensification, but future experiments should consider using real crop fields as the terrestrial matrix since they can represent different dispersal barriers.

## INTRODUCTION

Since the so-called Green Revolution, one of the cornerstones of industrial agriculture has been the management of agrochemicals such as pesticides, fertilizers, and soil amendment products (Foley et al., 2005; Schiesari et al., 2013; Schiesari & Grillitsch, 2011). These agrochemicals are applied in crop fields following specific schedules along the crop cycle (*e.g.* Schiesari et al., 2013), which is in itself frequently synchronized with the rainy season. Thus, not only direct overspray and drift but also runoff contributes to agrochemicals reaching surface water bodies (Carvalho, 2017; Matson et al., 1997). Despite the evident ecological importance of the seasonal and predictable release of these molecules, until recently, ecotoxicological studies were mostly devoid of ecological realism by emphasizing effects on individuals and populations instead of communities and ecosystems. Following this diagnosis, a relatively large number of studies mechanistically addressing direct and indirect effects of contaminants in local communities were conducted (*e.g.* Boone & James, 2003; Relyea et al., 2005; Rohr et al., 2006), while the importance of spatial context in modulating the effects of contaminants remains largely unexplored (Schiesari et al., 2018, 2019; but see Trekels et al., 2011). Yet, spatial structure is the norm in both natural and modified environments. For instance, many lentic aquatic environments can be found embedded in agricultural fields forming complex metacommunity structures (da Silva et al., 2012; Prado & Rossa‐Feres, 2014; Schiesari & Corrêa, 2016). Therefore, the consequences of agrochemicals and spatial structure on local communities must be addressed jointly if we are to manage biodiversity in agricultural landscapes successfully.

Agrochemical intensification can affect aquatic communities via two main pathways: First, it can positively or negatively affect primary productivity due to the use of organic and inorganic fertilizers, on one hand, and herbicides, on another (Isbell et al., 2013; Peterson et al., 1994; Rohr et al., 2006; Smith et al., 1999). Second, insecticides can strongly and negatively affect most aquatic insects, while indirectly benefiting non-insect taxa (Relyea, 2005a; Rohr et al., 2006). The use of NPK fertilizers is globally widespread (Foley et al., 2005), causing water nutrient enrichment, which leads to the increase of primary productivity and, consequently, algal biomass (Smith et al., 1999). Increased primary productivity can have several different consequences on higher trophic levels. For instance, it can increase competition, causing the abundance of consumers with a higher ability to deplete resources to increase (Abrams, 1988). In the presence of higher trophic levels, however, such effects can propagate through the trophic web via *bottom-up* effects, benefiting predators (Abrams, 1993; Cross et al., 2006; Slavik et al., 2004), which can then change consumer species composition by preferentially preying upon the most vulnerable prey (Leibold, 1996, 1999). Herbicides, on the other hand, can have opposite effects of fertilizers (Titeux et al., 2016). They are usually employed to control weeds in monocultures, but they can also affect non-target aquatic plants, such as green algae and macrophytes, possibly indirectly reversing the positive effects of nutrient enrichment on higher trophic levels (Rohr et al., 2006).

Insecticides have the potential to cause strong direct negative effects on any vulnerable species (Rohr et al., 2006). However, their indirect effects depend on the trophic level and competitive interactions among species that are differentially vulnerable to it. If the most vulnerable species are herbivores and detritivores, insecticides can indirectly and negatively affect predators, but positively affect producers. Alternatively, if the most vulnerable species are predators, herbivores and detritivores will be indirectly and positively affected by reduced predation, also negatively affecting producers (Relyea, 2005a; Relyea et al., 2005). The latter is the case of temporary freshwater pond systems, where aquatic insects are strongly negatively affected by pesticides, while their non-insect prey, amphibians, are not (Relyea, 2005a).

In a spatially structured landscape, dispersal can either mitigate or strengthen the effects of local selective pressures such as nutrient enrichment and pesticides (Leibold & Chase, 2018). For instance, if predators have a lower capacity to successfully colonize isolated habitats, bottom-up effects may be restricted to an increase in the abundance of herbivores and detritivores. This might be the case of aquatic insects, where predators have smaller populations and longer generation times than herbivores and detritivores; thus, they have fewer events of dispersal, which result in lower predator abundances in isolated habitats (Chase & Shulman, 2009; Kalinkat et al., 2015; Pelinson et al., 2020; Shulman & Chase, 2007). Additionally, because pesticides are usually applied in seasonal pulses, their direct acute effects can be, albeit strong, temporary. Therefore, their effects on communities can be understood as a temporary disturbance, followed by full or partial recovery of the original community structure (*i.e.*, Trekels et al., 2011). In this case, more isolated communities might take longer to recover from an insecticide pulse (Trekels et al., 2011).

In this study we aimed at understanding the consequences of chemical intensification on pond insect and amphibian community structure in different spatial contexts. We constructed replicated artificial ponds at different distances from a source wetland and experimentally simulated a gradient of chemical intensification by treating artificial ponds as if they were embedded in savannas (*i.e.*, no agrochemical use), pastures (*i.e.*, use of fertilizer), and sugarcane fields (*i.e.*, use of fertilizer and pesticides). We chose to simulate pastures and sugar cane fields because they are two of the most abundant land uses in Brazil, the largest country in the neotropical region and the most important sugarcane-producing region of the world (Zuurbier & van de Vooren, 2008). Pastures occupy around 20% of the Brazilian territory (~173 million hectares; MapBiomas, 2020), while sugarcane represents the third largest planted area in Brazil (~9 million hectares), only behind soybeans (~30 million hectares) and maize (~16 million hectares; IBGE, 2017). More importantly, because of the increasing demand for the replacement of fossil fuels for alternative biofuels (Titeux et al., 2016), sugarcane is predicted to expand in the next years, mostly over pasture and savanna areas (MapBiomas, 2020). We hypothesized that: (H1.1) Fertilization should change community structure in ‘pasture’ ponds by an increase in the abundance of predatory insects via bottom-up effects. (H1.2) This effect should be weaker in more isolated habitats where most predatory insects may not be able to establish large populations; thus, bottom-up effects should be stronger on herbivores and detritivores in more isolated ponds. (H2) The herbicide, on the other hand, should temporally reverse the effects of nutrient increase via a negative bottom-up effect in ‘sugarcane’ communities. (H3.1) We also expect that the insecticide will cause ‘sugarcane’ ponds to strongly diverge from other treatments, shifting community structure towards a higher abundance of amphibians due to a lower abundance of their predators. (H3.2) We further hypothesized that because predatory insects are predicted to be more dispersal limited than non-predatory ones, the possible positive indirect effects of the insecticide pulse on amphibians should last longer in more isolated habitats.

## METHODS

### Study area

We conducted a field experiment at the Estação Ecológica de Santa Bárbara (EESB) in Águas de Santa Bárbara, São Paulo, Brazil (22°48’59” S, 49°14’12” W). The EESB is a 2,712-ha protected area predominantly covered with open savanna phytophysiognomies, with smaller portions of seasonal semideciduous forests, *Pinus* sp., and *Eucalyptus* sp. plantations (Melo & Durigan, 2011). Soils are sandy and the climate is Koeppen’s Cwa, i.e., warm temperate with dry winters and hot summers (Kottek et al., 2006). Mean annual rainfall is ~1250mm with a distinct rainy season from October to March (January being the wettest month with ~200mm rainfall) and a dry season from April to September (July being the driest month with ~25mm rainfall; Hijmans et al., 2005). In the EESB, the experiment was implemented in an area covered by second-growth cerrado *sensu stricto*, a moderately dense, open-canopy savanna phytophysiognomy (Melo & Durigan, 2011).

### Experimental design

Experimental units consisted of ~1,200L artificial ponds dug into the ground and lined with a 0.5 mm thick, high-density polyethylene geomembrane to retain water. Each pond was 4m long, 1m wide, and 40 cm deep (Figure 1a). Walls were vertical along the length of the pond; 1m-long ramps terminating at ground level at each short side of the pond provided shallow microhabitats for freshwater organisms and escape for terrestrial fauna that eventually fell into the water (Figure 1a). Two roof tiles were placed at the waterline in each of the short sides to provide shelter and oviposition habitat for insects and amphibians. The experiment followed a fully factorial design crossing agrochemical intensification (simulating ‘savanna’, i.e., the control, ‘pasture’ and ‘sugar cane’ scenarios) with spatial isolation (three levels of isolation by distance from a putative source). Each land use-by-distance treatment was replicated five times for a total of 45 artificial ponds. The experiment ran from 19-Sep-2017 to 04-Mar-2018 and therefore encompassed the entire rainy season, effectively mimicking the dynamics of temporary ponds that are common in both preserved and converted landscapes. Between 19 and 25-Sep-2017, mesocosms were filled with well water. On 30-Sep-2017, we added to each mesocosm 800g (wet mass) of leaf litter composed of equal amounts of grass and tree leaf litter to provide a source of nutrients and structural complexity for benthic organisms.

**Figure. 1.**
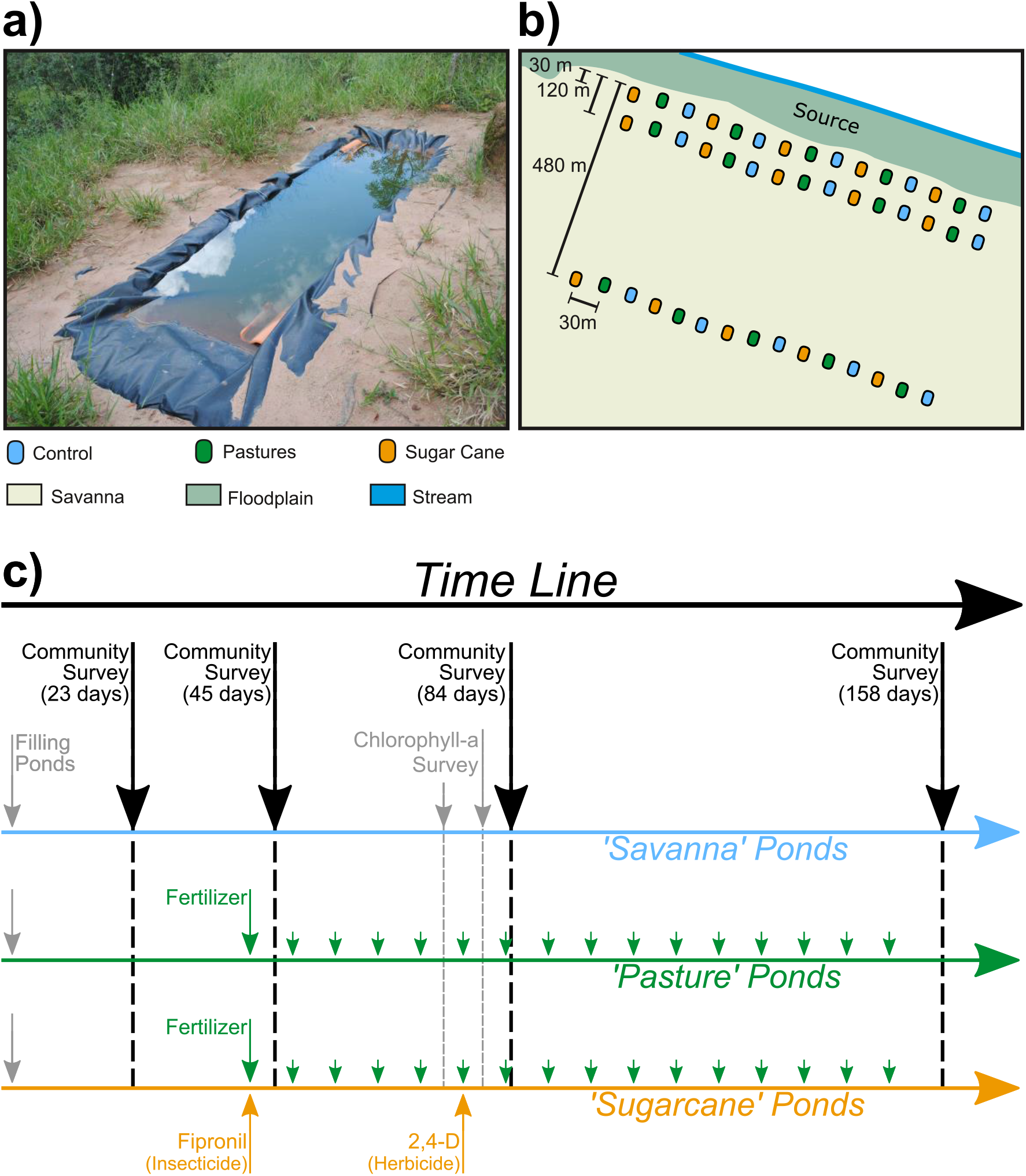
(a) One of the naturally colonized pond mesocosms during the experiment. (b) Experimental setup comprising three rows of 15 artificial ponds at 30, 120, and 480 m from a source water body (a stream and its floodplain). (c) Temporal design comprising pond filling; community surveys after 23, 45, 84, and 158 days; and agrochemical management according to the ‘crop cycle’ approach.

### Spatial Isolation

The spatial isolation treatments were achieved by establishing 15 artificial ponds along each of three parallel transects 30m, 120m, and 480m from a source wetland consisting of a stream (Riacho Passarinho) and its floodplain (Figure 1b). Within each transect, the distance between adjacent artificial ponds was 30 m. The well-drained sandy soils ensured that no other ponds and puddles formed during the rainy season at our study site, which could confound our manipulation of isolation distances. Ponds were naturally colonized by aquatic insects and amphibians throughout the rainy season. The chosen isolation distances represent a steep (X16) gradient known to produce important differences in abundance patterns for the majority of the taxa that we expected to colonize the artificial ponds (*e.g.* Chase & Shulman, 2009; Pelinson et al., 2020; Shulman & Chase, 2007; Silva et al., 2012; Trekels et al., 2011; Wilcox, 2001).

### Land use treatments

Our experiment followed a ‘crop cycle’ design (Arts et al., 2006; van Wijngaarden et al., 2004) in that molecules, doses, and schedule of application were realistically simulated. The ‘pasture’ agrochemical treatment was achieved by weekly nutrient additions, whereas the ‘sugarcane’ treatment was achieved by identical weekly nutrient additions plus one insecticide and one herbicide application pulses (Figure 1c). For representativeness, we simulated conventional sugarcane land management practices in Central-South Brazil at the ratoon cropping stage (the sugarcane cycle typically lasts 5 to 6 years with yearly harvests via ratoon cropping; Therefore, at any given moment 4/5 to 5/6 of all sugarcane fields are at a ratoon cropping stage; see supplement 1 for details).

The manipulated commercial formulation NPK formulation was composed of ammonium nitrate (NH_4_NO_3_), monoammonium phosphate (NH_4_H_2_PO_4_), and potassium chloride (KCl) following a 20-05-20 NPK ratio (20% of N, 5% of P_2_O_5_, and 20% of K_2_O) at a dose of 80 kg/ha of N, 20 kg/ha of P_2_O_5_ and 80 kg/ha of K_2_O. This dosage is compatible with the fertilization of both moderately managed pastures (Santos et al., 2010) and ratoon cane (Cantarella & Rossetto, 2010). Dosage was scaled to pond surface area (4 m², *i.e.*, 4 × 10^−4^ ha) and divided into equal weekly doses to simulate a linear dissolution over 52 weeks (*i.e.*, one year), which is the usual interval for fertilizer reapplications. An amount of 3.08 g of the NPK 20-05-20 formulation was added weekly from 30-Oct-2017 (the simulated planting or harvesting date, when fertilizers are typically applied) until the end of the experiment, for a total of 15 applications. Therefore, by the end of the experiment, each ‘pasture’ and ‘sugarcane’ pond had received a total of 46.2 g of NPK 20-05-20.

Commercial pesticides selected for manipulation were REGENT ® 800 WG (BASF; active ingredient fipronil at a concentration of 800 g AI/kg), and DMA ® 806 BR (Dow AgroSciences. active ingredient 2,4-D at a concentration of 806 g AI/L). These active ingredients are among the top-selling insecticides and herbicides employed in sugarcane plantations in the State of São Paulo (see details in supplement 1). Pesticides were applied at the maximum doses recommended by the manufacturer for pest and post-emergence weed control in sugarcane plantations at the ratoon cropping stage (250 g/ha of REGENT ® 800 WG; 1,5 L/ha of DMA ® 806 BR). As above, doses were scaled to pond surface area. We made one single application pulse of the insecticide on 30-Oct-2017 (the simulated planting or harvesting date) and one single application pulse of the herbicide five weeks later, on 04-Dec-2017, which is a common timeframe for the application of post-emergence herbicides (R. Rossetto *pers. com*). Pesticide applications were made by diluting 0.1 g of REGENT ® 800 WG (*i.e.*, 80 mg of fipronil) or 0.6 ml of DMA ® 806 BR (*i.e.*, 483,6 mg of 2,4-D) in 400 ml of well water and applying this solution directly into each ‘sugarcane’ pond. After each application, we gently steered the water with a wood stick and collected a 20 ml subsurface water sample of each of the 15 ‘sugarcane’ ponds to produce a 300 ml composite sample that was analyzed for pesticide concentrations. We did not attempt to thoroughly mix mesocosms to avoid mechanical damage to the established community, which was to be sampled a few days later. One last composite water sample was also collected at the end of the experiment. Water samples were stored in ice, protected from light, and transported to the lab for pesticide dosage within one day. Measured concentrations at the time of application were 15.6 μg L^−1^ for fipronil and 337.8 μg L^−1^ for 2,4-D. Pesticide concentrations were below detection and quantification limits by the end of the experiment for fipronil and 2,4-D respectively (see supplement 2 for details). We also measured total nitrogen (TN) and total phosphorus (TP) (see supplement 3 for details).

### Community survey

Because community composition changes drastically in the earliest phases of colonization, we proposed to conduct community surveys 20, 40, 80, and 160 days from the start of the experiment. Actual sampling dates were 13 to 19-Oct-2017, 03 to 10-Nov-2017, 12 to 19-Dec-2017, and 24-Feb-2018 to 03-Mar-2018 (about 23, 45, 84, and 158 days from the start of the experiment). Because the 23-days survey was taken before agrochemical additions, here we will only show results for the three last surveys (but see supplement 4 for results of the 23 days survey). Samples were taken by pipe sampling, which provided quantitative per-unit-area information on species composition and abundances. The sample was taken by quickly thrusting the pipe through the water column and into the sediments to seal the sample area. We then swept all the pipe bottom (area 0,102 m²) four times and the pelagic area three times with a hand net (mesh size 1.5 mm). We took four samples, as described above, per pond, which were summed, at each sampling survey. Therefore, we ended up with one sample unit per pond per survey. After samples were cleaned of sediment and debris, tadpoles were immediately euthanized and preserved in 10% buffered formalin and invertebrates in 70% ethanol. We counted and identified all aquatic macroinvertebrates to the genus level, with an exception for the Chironomidae and Ceratopogonidae families, which were identified to subfamily and family levels, respectively. Amphibians were assigned to species level, except for *Scinax*, which were identified to the genus level.

### Chlorophyll-a survey

Because our experiment aimed at observing the net effects of chemical intensification simulating a full crop cycle, we could not assess the separated effects of insecticides and herbicides on community structure. Therefore, to better assess the possible effects of the herbicide application, we also sampled the *in vivo* chlorophyll-a concentration in water as a surrogate for algae biomass responses to the herbicide application. Chlorophyll-a concentration was assessed before (02-Dec-2017) and after (08-Dec-2017) herbicide application (04-Dec-2017) by fluorescence with a portable fluorometer (Aquafluor, Turner Designs, San Jose, CA, USA).

### Data Analysis

#### Total abundances and Chlorophyll-a

We tested whether land-use treatments, spatial isolation, sampling survey, and their interaction caused any changes in total abundance of predatory insects, non-predatory insects, amphibians, and chlorophyll-a biomass separately through type II Wald chi-square tests. We considered predatory insects only the predators that were prone to prey upon the other sampled taxa. Insects that are not predators, that prey mostly upon zooplankton, or that have only a small portion of their diet based on predation were considered non-predatory insects (*i.e.* herbivores and detritivores; see supplement 5). We modeled total abundances and chlorophyll-a concentration using generalized linear mixed models (GLMMs) with negative binomial and gamma distributions, respectively, and pond identifications as a random effect term. We also performed pairwise post-hoc tests with Tukey adjustment of p-values for factors that significantly affected total abundance patterns.

#### Community structure

To test the hypothesis that community structure is influenced by the agrochemical treatments, distance to the source, and their interaction, we used a generalized linear model approach for multivariate data where the matrix of site-by-raw species abundance data represents community structures (Warton et al., 2015). Before analyzing our community data, we removed all species with three or fewer occurrences in each survey, both because they are uninformative to general community patterns and because they complicate model parameter estimation (Warton et al., 2015).

Because multivariate patterns can be relatively complex and the effect of treatments are expected to change over time, we analyzed data from each survey separately. Because species abundances were overdispersed, we modeled them with a negative binomial distribution. To test whether different treatments and their interaction had a significant effect on community structure, we performed likelihood ratio tests between models with and without each model term but keeping all other lower-order terms (*i.e.* type II test). To account for the non-independence between species abundances when computing p-values, we used a permutation approach that shuffled entire rows of the incidence matrix (ponds), keeping species abundances in the same ponds always together. P-values were computed using the PIT-trap bootstrap resample procedure from 1,000 bootstrap resamples, which operate on probability integral transform residuals (Warton et al., 2017). We also performed pairwise post-hoc tests with false discovery ratio (FDR) correction of p-values for factors that significantly affected community structure patterns. To better interpret changes in community structure between treatments in terms of the groups we previously defined (*i.e.* predatory insects; non-predatory insects and amphibians), we refitted the models with the significant predictor terms adding species functional groups as a predictor of the community responses to treatments. This approach is called the model-based fourth corner solution (Brown et al., 2014). To assess whether individual taxa or functional groups significantly respond to the different treatments, we assessed 95% confidence intervals of the pairwise differences in abundances between treatments. Note that this approach differs from looking at total abundances within functional groups because it assumes equal weights to different taxa within each functional group, regardless of its total abundance. Those analyses were implemented using functions *manyglm(), anova.manyglm()*, and *traitglm()* from package ‘mvabund’ version 4.0.1 (Wang et al., 2012, 2020).

To visualize multivariate differences in community structure, we performed an unconstrained ordination for each of the surveys separately using generalized linear latent variable models (GLLVM) with two latent variables (Hui et al., 2015; Niku et al., 2017). The latent variables were estimated via variational approximation (Hui et al., 2017). These analyses were implemented using the function *gllvm()* from package ‘gllvm’ version 1.1.3 (Niku et al., 2019). All analyses were implemented in software R version 3.6.1 (R Core Team, 2020).

## RESULTS

### Total abundance and Chlorophyll-a

Mesocosms were colonized by amphibians, and aquatic and semiaquatic insects comprising six orders and 22 families (see supplement 5 for details). The abundance of predatory insects, non-predatory insects, and amphibians were all affected by time since the beginning of the experiment (Table 1). The total abundance of predatory insects and amphibians was highest at the 84-days survey (*i.e.* middle of the rainy season) (Figure 2a and 2c), whereas non-predatory insects were more abundant by the end of the rainy season (*i.e.* 158-days survey) (Figure 2b).

**Table 1.** Summary of deviance analysis (type II Wald Chi-square tests) for total abundances of predators, non-predatory insects, amphibians, and chlorophyll-a concentration. Significant terms are highlighted in bold (p < 0.05).

|  | Degrees of freedom | Chisq | p |
| --- | --- | --- | --- |
| *Predatory insects* |  |  |  |
| **Land Use** | **2** | **19.286** | **<0.001** |
| Isolation | 2 | 0.385 | 0.825 |
| **Survey** | **2** | **89.563** | **<0.001** |
| Land Use : Isolation | 4 | 5.356 | 0.253 |
| **Land Use : Survey** | **4** | **39.955** | **<0.001** |
| Isolation : Survey | 4 | 7.425 | 0.115 |
| Land Use : Isolation : Survey | 8 | 14.166 | 0.078 |
| *Non-predatory insects* |  |  |  |
| **Land Use** | **9** | **114.588** | **<0.001** |
| **Isolation** | **7** | **47.363** | **<0.001** |
| **Survey** | **7** | **230.224** | **<0.001** |
| **Land Use : Isolation** | **7** | **16.550** | **0.021** |
| **Land Use : Survey** | **7** | **95.333** | **<0.001** |
| **Isolation : Survey** | **7** | **25.160** | **0.001** |
| **Land Use : Isolation : Survey** | **8** | **21.912** | **0.005** |
| *Amphibians* |  |  |  |
| Land Use | 2 | 2.408 | 0.300 |
| **Isolation** | **2** | **9.657** | **0.008** |
| **Survey** | **2** | **39.887** | **<0.001** |
| Land Use : Isolation | 4 | 1.593 | 0.810 |
| **Land Use : Survey** | **4** | **9.451** | **0.051** |
| **Isolation : Survey** | **4** | **9.837** | **0.043** |
| Land Use : Isolation : Survey | 8 | 7.182 | 0.517 |
| *Chlorophyll-a* |  |  |  |
| **Land Use** | **2** | **13.289** | **0.001** |
| Isolation | 2 | 1.322 | 0.516 |
| **Time (relative to herbicide application)** | **1** | **8.882** | **0.003** |
| Land Use : Isolation | 4 | 3.134 | 0.536 |
| Land Use : Time | 2 | 2.192 | 0.334 |
| Isolation : Time | 2 | 0.407 | 0.816 |
| Land Use : Isolation : Time | 4 | 6.638 | 0.156 |

**Figure. 2.**
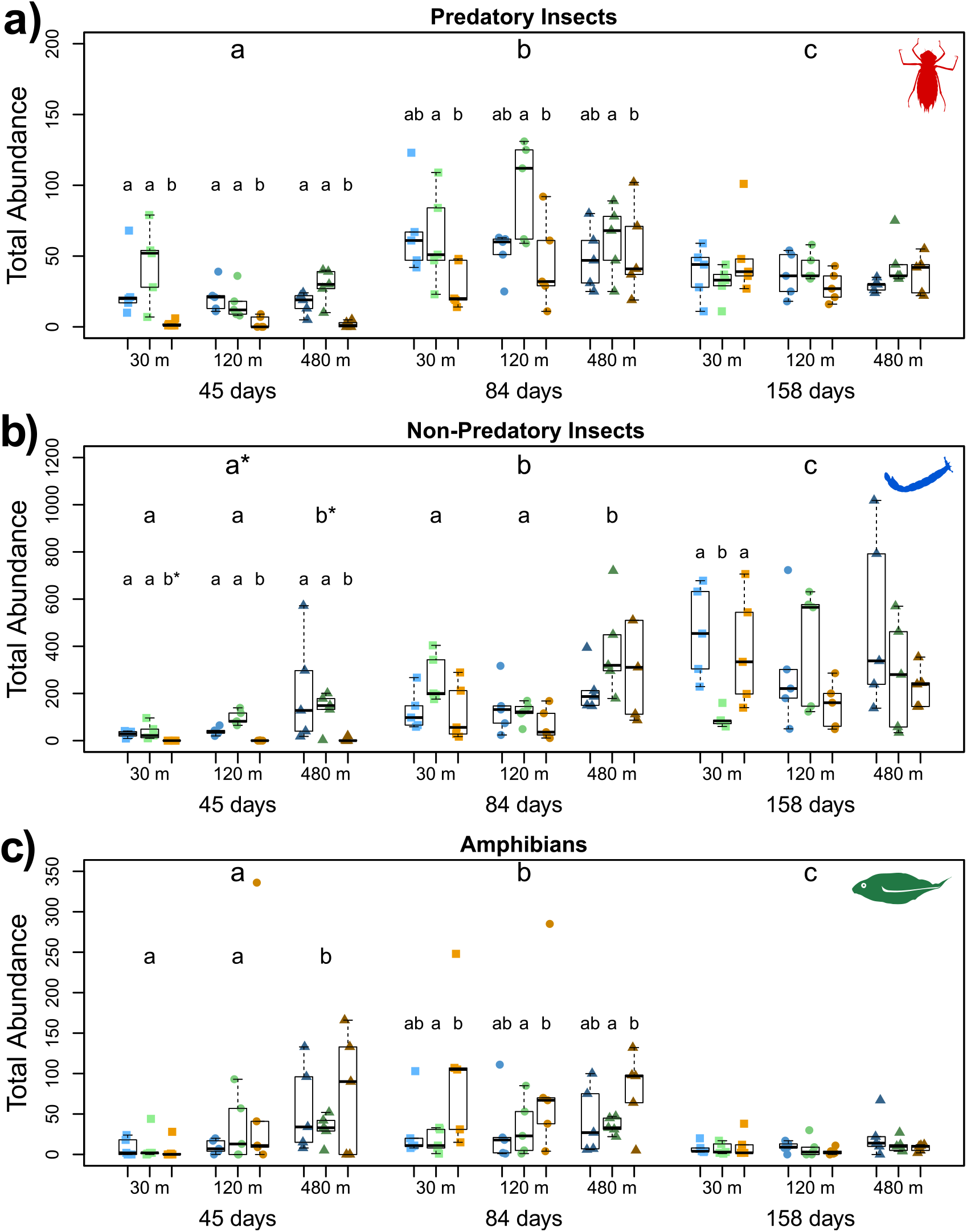
Total abundance of predatory insects (a), non-predatory insects (b), and amphibians (c) across the different surveys, isolation, and land use treatments. Blue, green and yellow symbols are ‘savanna’, ‘pasture’, and ‘sugarcane’ ponds, respectively. Squares, circles, and triangles are 30 m, 120 m, and 480 m isolation treatments. Different letters represent statistical differences among treatments. Pairwise comparisons were done across surveys in all cases (a, b, and c). When there was a significant interaction between the survey and isolation, pairwise comparisons were done between isolation treatments in each survey separately (b and c); when there was a significant interaction between land use and survey, they were done between land use treatments for each survey separately (c); when there was a significant interaction between survey, land use, and isolation, pairwise comparisons were done between land use treatments for each survey and isolation treatment separately (a).

Spatial isolation did not affect the abundance of predatory insects (Table 1; Figure 2a), however, non-predatory insects were more abundant in the 480 m ponds at both the 45- and 84-days surveys (Table 1; Figure 2b). Amphibians were also more abundant in the 480 m ponds, but only at the 45-days survey (Table 1; Figure 2c).

The abundance of both predatory and non-predatory aquatic insects was strongly reduced in ‘sugarcane’ ponds at the 45-days survey (*i.e.* right after insecticide application) (Table 1; Figures 1a and 1b). Predatory insects were still less abundant in ‘sugarcane’ ponds at the 84-days survey if compared to pasture treatments (Figure 2a). The total abundance of amphibians only exhibited a marginally significant interaction term between land use treatment and survey (Table 1). They were more abundant in ‘sugarcane’ ponds at the 84-day survey, but only when compared to ‘pasture’ ponds (Figure 2c). The total abundance of predatory insects was higher in ‘pasture’ ponds in the middle of the rainy season, whereas amphibians exhibited the opposite pattern (Figures 2a and 2c).

Chlorophyll-a concentration, contrary to our expectation, was higher in all treatments after ‘sugarcane’ ponds were treated with the herbicide (Table 1; Figure 3). It was also generally higher in ‘sugarcane’ ponds if compared to ‘savanna’ treatments (p = 0.002).

**Figure 3.**
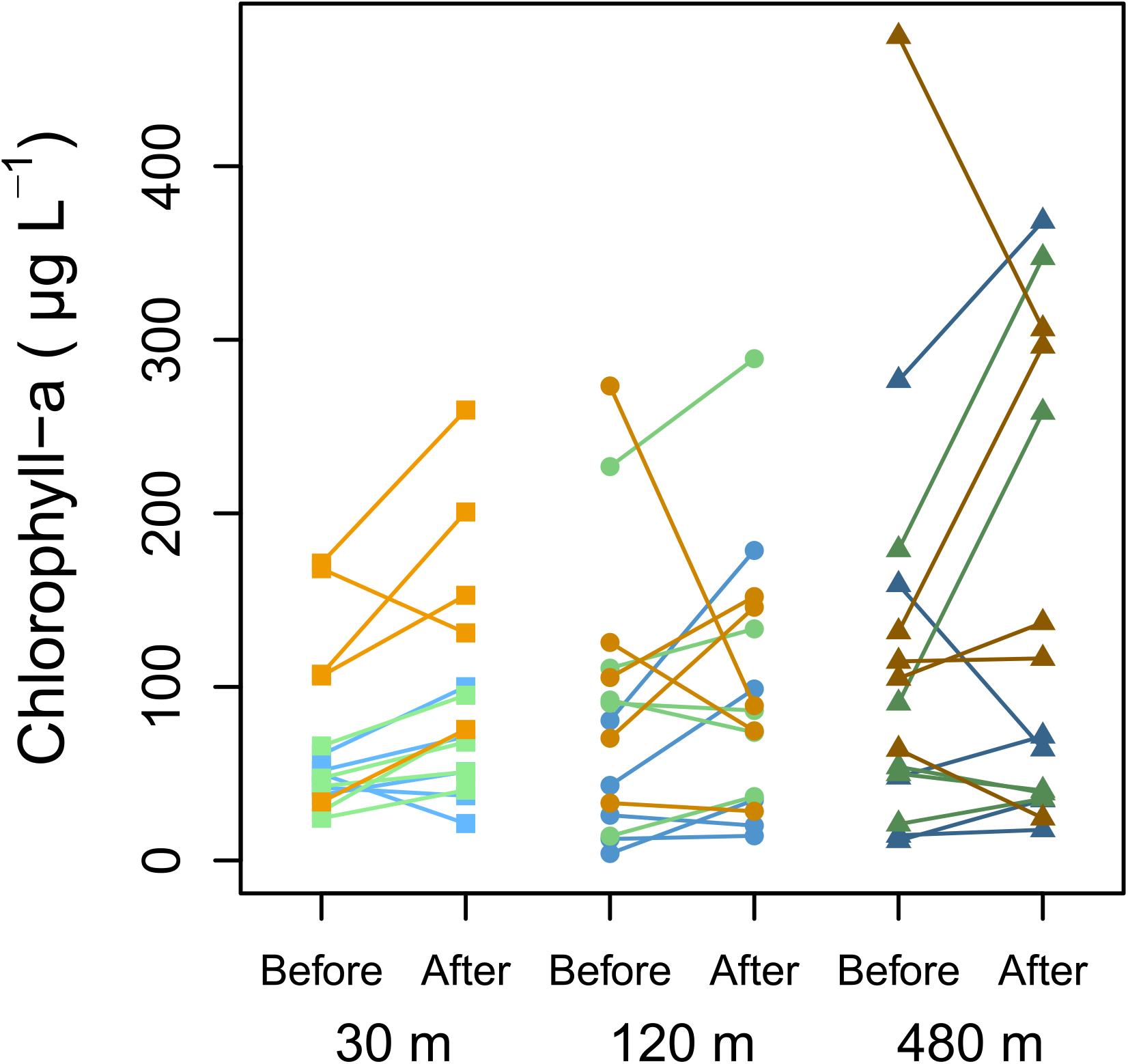
Chlorophyll-a concentration before and after the herbicide application (only in ‘sugarcane’ ponds) in each isolation distance separately. Blue, green, and yellow symbols are ‘savanna’, ‘pasture’, and ‘sugarcane’ ponds respectively. Symbols representing the same ponds before and after the herbicide application are connected by a line.

### Community structure

We did not find interactive effects among land use and spatial isolation treatments, but both factors affected community structure across the entire rainy season (Table 2; Figure 4). Ponds at 30 m of isolation were never significantly different from those at 120 m, but the 30 m distance ponds were always different from ponds at 480 m. Ponds at 120 m were only different from those at 480 m at the 84- and 158-days surveys. At the 45-days survey, these differences were driven by general increases in the abundances of amphibians and non-predatory insects in more isolated ponds (Figure 5a). At 84-days, it was mostly driven by increases in the abundance of non-predatory insects in more isolated ponds and idiosyncratic responses of predatory insects to isolation (see supplement 6). At 158 days, these patterns were driven by negative responses of both predatory and non-predatory insects to isolation (Figure 5a).

**Table 2.**
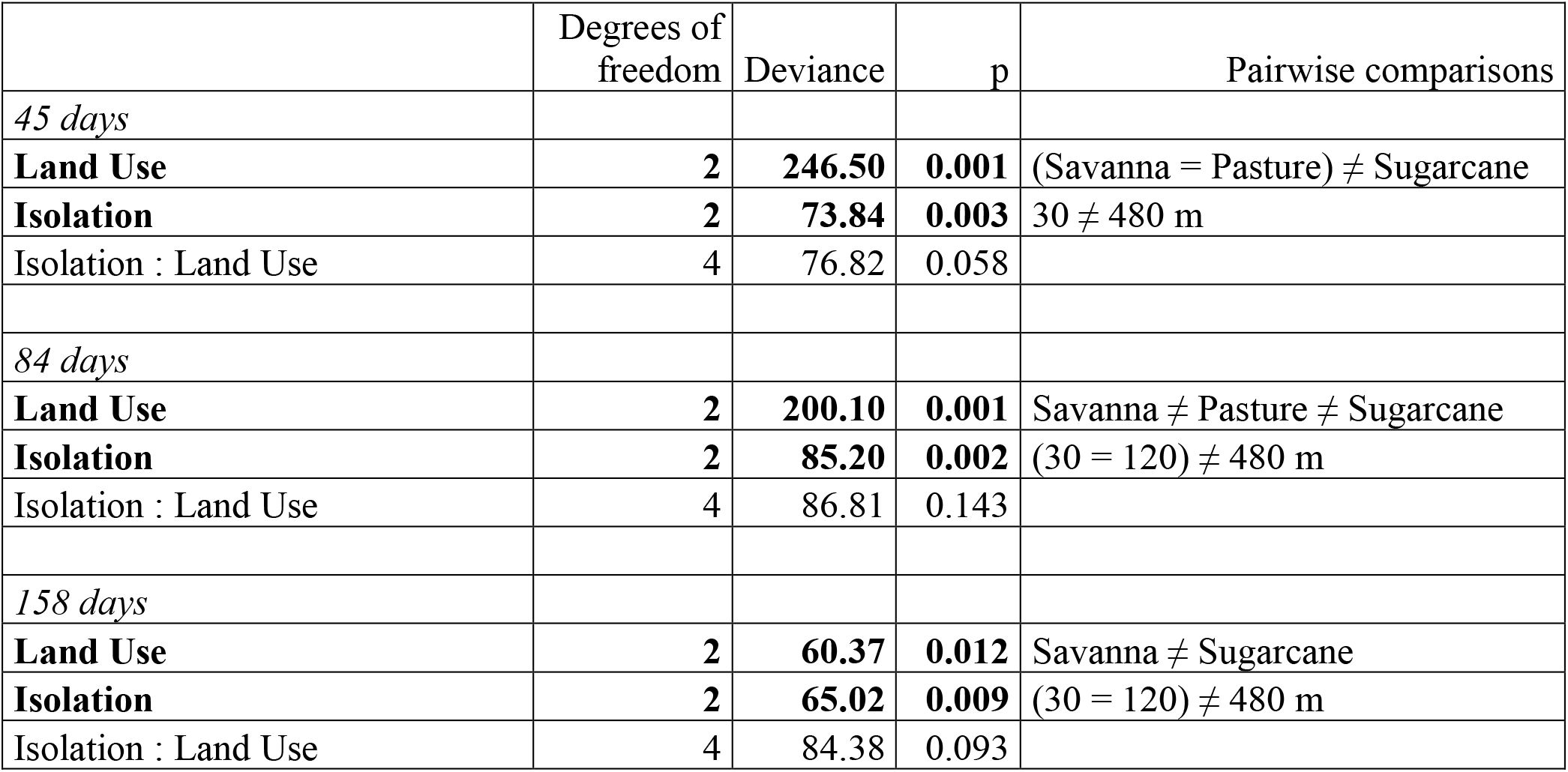
Summary of deviance analysis (likelihood ratio tests) for community structure in each survey separately. Significant terms are highlighted in bold (p < 0.05). Only significant pairwise differences are shown.

|  | Degrees of freedom | Deviance | p | Pairwise comparisons |
| --- | --- | --- | --- | --- |
| *45 days* |  |  |  |  |
| **Land Use** | **2** | **246.50** | **0.001** | (Savanna = Pasture) ≠ Sugarcane |
| **Isolation** | **2** | **73.84** | **0.003** | 30 ≠ 480 m |
| Isolation : Land Use | 4 | 76.82 | 0.058 |  |
| *84 days* |  |  |  |  |
| **Land Use** | **2** | **200.10** | **0.001** | Savanna ≠ Pasture ≠ Sugarcane |
| **Isolation** | **2** | **85.20** | **0.002** | (30 = 120) ≠ 480 m |
| Isolation : Land Use | 4 | 86.81 | 0.143 |  |
| *158 days* |  |  |  |  |
| **Land Use** | **2** | **60.37** | **0.012** | Savanna ≠ Sugarcane |
| **Isolation** | **2** | **65.02** | **0.009** | (30 = 120) ≠ 480 m |
| Isolation : Land Use | 4 | 84.38 | 0.093 |  |

**Figure 4.**
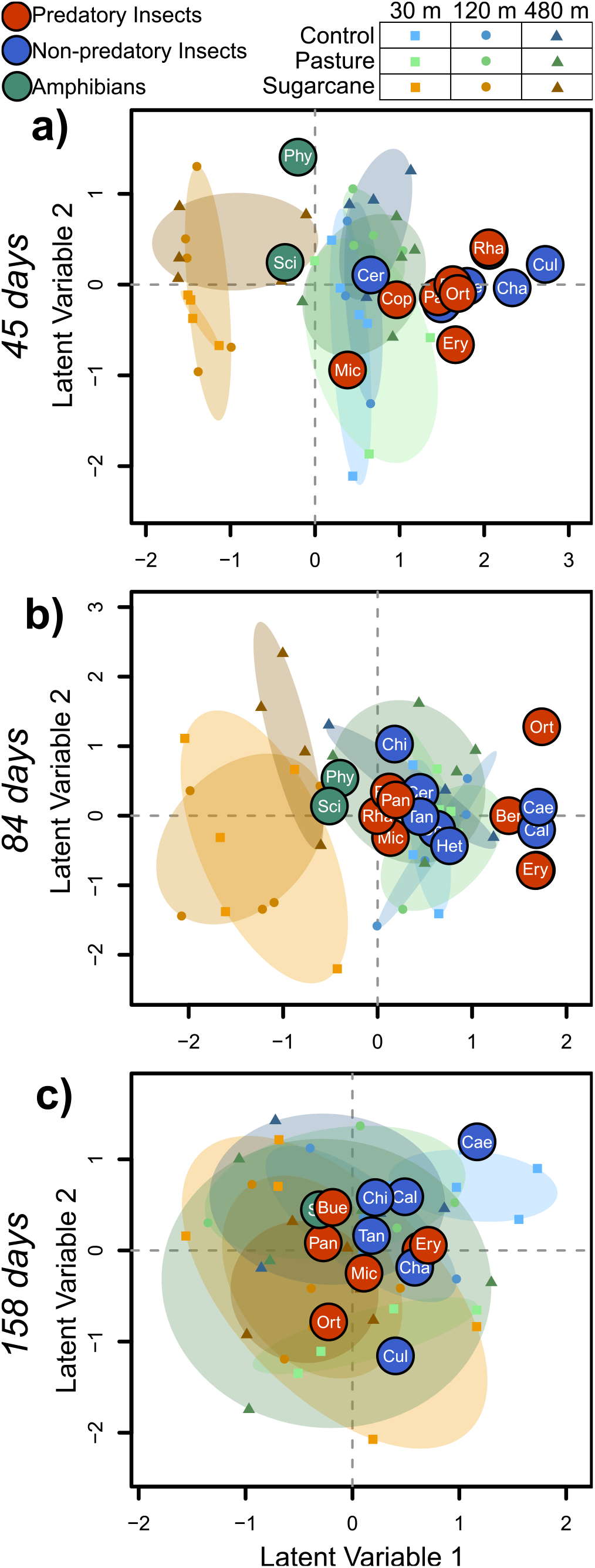
Model-based unconstrained ordinations showing pond communities (symbols) and species (bubbles) in each of the community surveys (a to c). Abbreviations of names of taxa are provided in supplement 5.

**Figure 5.**
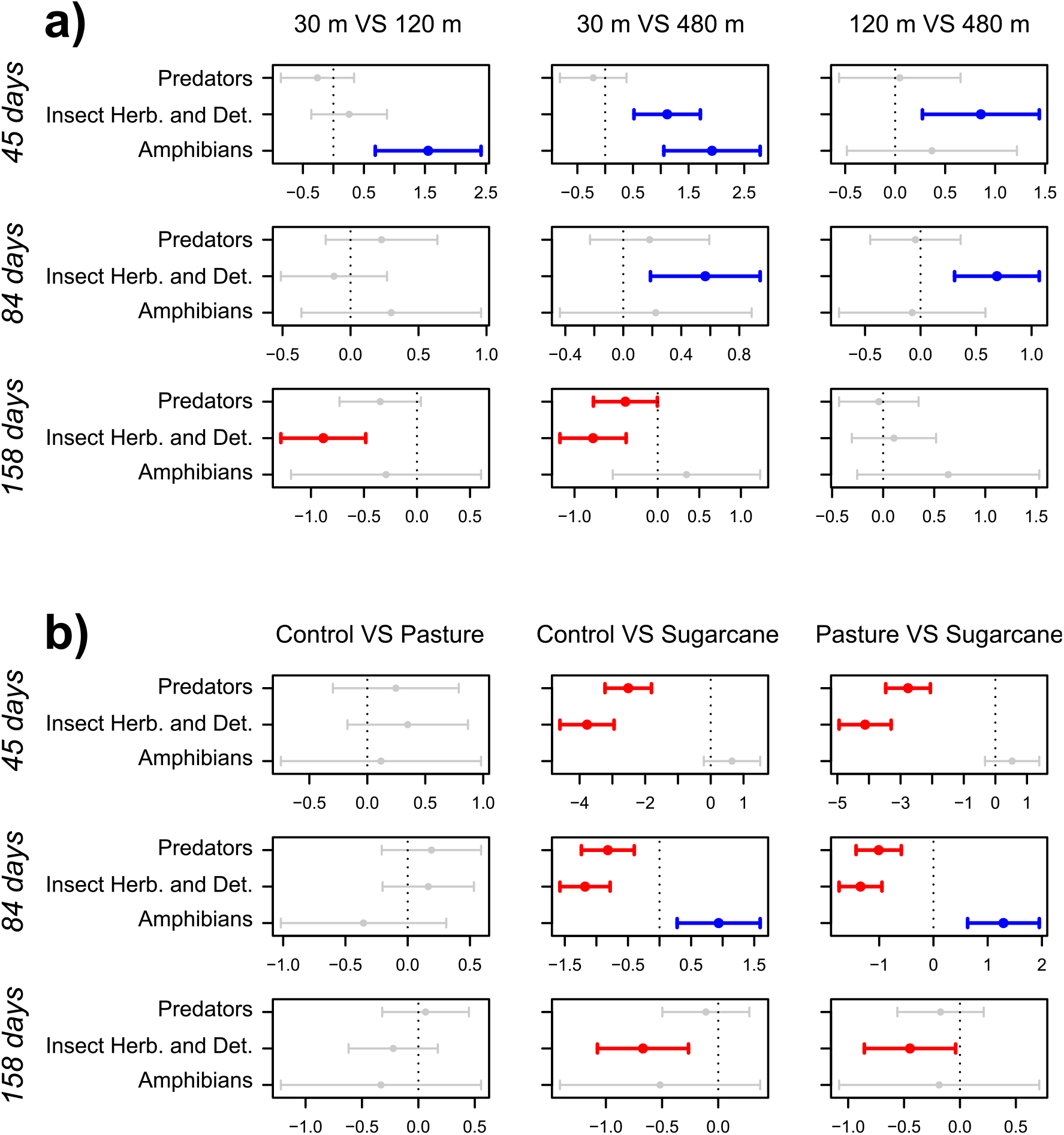
Pairwise maximum likelihood estimates (dots) and their 95% Confidence intervals (bars) of the differences between isolation (a) and land use (b) treatments for each functional group. Confidence intervals not crossing the zero hatched line were considered significant effects and colored; blue bars represent an increase and red bars a decrease in abundance from the reference treatment (reference VS treatment). These are the mean effects of treatments on different groups of taxa (*i.e.* predatory insects, non-predatory insects, and amphibians). Similar estimates for individual taxa are presented in supplement 6.

Community patterns regarding land use treatments were dominated by differences of ‘sugarcane’ to ‘savanna’ and ‘pasture’ communities after the insecticide application throughout the entire experiment (Figure 4). ‘Savanna’ ponds were always different from ‘sugarcane’ ponds (Table 2). At the 45-days survey, these differences were driven by reductions in the abundance of most aquatic insects whereas amphibian populations persisted in all treatments (Figures 4a and 5b). At the 84-days survey, differences were still driven by reductions in the abundances of aquatic insects, but also an increase in the abundances of amphibians in ‘sugarcane’ ponds. At the 158-days survey, differences were still driven by reductions in the abundances of non-predatory insects, but predators showed both positive and negative responses (see supplement 6). At this stage of community assembly, predatory insects and amphibians had similar abundances in both treatments (Figure 5b). Differences between pasture and sugarcane treatments were similar to those between ‘savanna’ and sugarcane treatments, but milder at the 158-days survey (Table 1; Figure 5b). ‘Pasture’ ponds were only significantly different from ‘savanna’ ponds at the 84-days survey (Table 2), however, this difference was driven by just idiosyncratic changes in the abundance of a few predatory insects (see supplement 6).

## DISCUSSION

We were able to assess how temporary pond communities change as a consequence of chemical intensification in different spatial contexts by following community assembly for an entire rainy season, which coincides with the application of fertilizers and pesticides in real moderate-intensity pastures and sugarcane crop fields. We saw that ‘pasture’ communities were only slightly different from ‘savanna’ communities in the middle of the rainy season. ‘Sugarcane’ communities, however, largely diverged from ‘savanna’ and ‘pasture’ communities, even though this effect lost strength with time. Spatial isolation showed a generally positive effect on non-predatory insects, as expected, but only idiosyncratic effects upon predatory ones. Amphibians were surprisingly only positively affected by isolation. Furthermore, we found no evidence that isolation by distance can cause important changes in the effects of agrochemicals on freshwater insect and amphibian communities.

Spatial isolation positively affected the total abundance of non-predatory insects, consequently affecting community patterns. We expected that such positive responses would be an indirect consequence of a potential reduction of predation in isolated ponds. However, contrary to our expectations, we did not observe a reduction in the total abundance of predatory insects across the gradient of spatial isolation. We believe there may be two possible non-mutually exclusive explanations for these patterns. First, the few predatory insects that were negatively affected by isolation (*e.g. Erythrodiplax* dragonflies and *Berosus* beetles; see supplement 6) could be the ones that preferentially prey upon most non-predatory insects (*e.g.* Chironomid midges, *Culex* mosquitoes; see supplement 6). Second, the total abundance patterns exhibited by predatory insects could be a net consequence of both a negative effect of isolation and a positive effect of higher prey availability in more isolated habitats, especially amphibians, which were highly abundant in isolated ponds since the very beginning of the experiment (see supplement 4). Indeed, amphibians were absent from an analogous experiment we conducted previously, likely because it started late in the rainy season. In that case, even though predatory insects exhibit idiosyncratic responses to isolation by distance, their total abundance was generally reduced (see Pelinson et al., 2020).

Amphibians exhibited no negative effects of isolation at any moment during the entire rainy season. This is surprising given the overwhelming evidence that amphibians have limited dispersal ability due to their high risk of desiccation during terrestrial dispersal (*e.g.*, Marsh et al., 1999; Marsh & Trenham, 2001; Rittenhouse & Semlitsch, 2007; Semlitsch, 2008; Sinsch et al., 2012). We believe that there could be two joint explanations for such a pattern. First, habitat selection by amphibians is subject to context dependence (Resetarits, 2005), where adults should perceive habitat quality differently depending on the surrounding context (see Resetarits & Silberbush, 2016). Specifically, amphibians are known to strongly avoid habitats containing fish (Resetarits, 2005; Resetarits & Wilbur, 1991; Vonesh et al., 2009). In this case, ponds that are closer to the source wetland (*i.e.* the stream and its marshy floodplains) could be perceived as having a comparatively lower quality (*i.e.*, higher risk of predation by fish), whereas isolated ponds would be perceived as high-quality habitats since they are far away from the source. Second, because we conducted our experiment in a relatively dense savannah landscape with sparse trees, the terrestrial matrix could have been a suitable habitat for amphibian adults to use for shelter and foraging, especially for generalist amphibians with adaptations to living and reproduce in dryer habitats, as the species that colonized our mesocosms are (Haddad & Prado, 2005; Vasconcelos et al., 2014). It would mean that these amphibian species could have been uniformly distributed across the terrestrial habitat, regardless of the distance from the source. It is important to mention that this might be an important deviation between our manipulated environmental scenario and those observed in true agricultural landscapes. For instance, Silva et al. (2012) found a strong reduction in adult anuran abundance in pastures after 50 m of distance from forest fragments. True open canopy agricultural landscapes might vary in permeability to dispersal (*e.g.*, Hansen et al., 2019)), with open canopy ones, such as pastures, being likely drier and more hostile habitats to dispersal if compared to dense savannahs or forest fragments (*e.g.*, Rothermel & Semlitsch, 2002, 2002; Watling & Braga, 2015). Therefore, future field experiments that consider the spatial context of aquatic communities should consider using real crop fields as their terrestrial matrix.

Because the ‘pasture’ ponds were only treated with fertilizers, we expected a general increase in the abundance of predatory insects (H1.1; Abrams, 1993; Slavik et al., 2004). We indeed observed that the total abundance of predatory insects was slightly higher in ‘pasture’ ponds in the middle of the rainy season (*i.e.* 84-days survey). When we looked at community structure patterns, however, the small differences among ‘pasture’ and ‘savanna’ ponds were not driven by consistent increases in predatory insect abundances. Digging deeper into community patterns, only *Orthemis* dragonflies and *Microvelia* positively responded to pasture treatments in the middle of the rainy season (see supplement 6). Similar increases in the abundance of predators have been observed before. For instance, Jahnke et al. (2001) found that the majority of the taxa positively correlated with a gradient of nutrient concentration in wetlands were predacious dytiscid beetles. Still, even though noticeable, the increase in abundance of both these predatory taxa, and of all predatory taxa together, was mild, even though we had already added a large amount of the NPK fertilizer to each pasture and ‘sugarcane’ pond (roughly about 6.3 mg L^−1^ of nitrogen and 3.6 mg L^−1^ of phosphorus). We believe that the differences we found were small because these temporary ponds are often in the hypertrophic portion of the primary productivity gradient. For instance, (Schiesari & Corrêa, 2016) observed that ponds embedded in either savannah, pasture, or sugarcane landscapes had values of TP and TN above 1000 ug L^−1^, which are way higher than expected for eutrophic lakes (*i.e.*, between 30 and 100 ug L^−1^, Smith et al., 1999). Indeed, the water surface levels of TP and TN measured in the middle of the rainy season in both pasture and ‘savanna’ ponds were not significantly different and almost always above 100 ug L^−1^ (see Supplement 3). Still, even in this extreme of the gradient, we were able to observe that intensification of pastures by using fertilizers can potentially affect aquatic communities, likely through bottom-up effects.

We did not find differences in the effect of the fertilization regime in ‘pasture’ ponds in different spatial isolation treatments (H1.2). We expected that, because predatory insects are usually more abundant in less isolated ponds, the positive *bottom-up* effect of fertilization would be stronger upon predatory insects in less isolated ponds, and upon herbivores and detritivores in more isolated ones. The lack of this interactive effect may have happened because, as we already discussed, predatory insects were not consistently negatively affected by isolation, thus, any increase in abundance of non-predatory taxa in more isolated ponds would have been buffered by predator consumption via *top-down* regulation (Abrams, 1993), just as we expected for less isolated ponds.

We did not observe any clear acute negative effects of the herbicide 2,4-D on species abundance patterns after the application (H2), which is not surprising. Relyea (2005) also found no differences in general abundance patterns of predatory insects, amphibians, or snail herbivores that were treated with similar concentrations of 2,4-D. These results are also consistent with past toxicity studies with amphibians (LC50_96h_ varying from 28.8 to 574.2 mg L^−1^, depending on the species, (Freitas et al., 2019) and aquatic insects such as chironomids (Pinto et al 2001), which only show no acute effects with concentrations similar to what we manipulated. We also did not observe differences in the response of chlorophyll-a biomass to herbicide application. This is also consistent with past studies that show that 2,4-D has moderate toxicity to macrophytes but low toxicity to algae (Peterson et al., 1994; PPDB, 2020, but see Moreira et al., 2020), which significantly decreases its potential to cause negative *bottom-up* effects (H2) in temporary ponds in general. However, the non-lethal effects of 2,4-D could still have played an important role in our experiment. 2,4-D has been found to decrease the swimming speed of tadpoles at the same concentrations that we manipulated (Freitas et al., 2019), possibly making amphibians more susceptible to predation. However, we do not know what systemic effects 2,4-D could have on predatory insects. For instance, other pesticides have been found to increase prey survival in the presence of predators because it decreases their predatory potential (Hanlon & Relyea, 2013). It is also important to mention that, even though we found no evidence that 2,4-D can cause significant changes in insect and amphibian communities, many works have shown that glyphosate-based herbicides (top-selling herbicide in the state of São Paulo) that contain the surfactant polyethoxylated tallowamine in their formulation, such as Roundup ®, are highly toxic to amphibians (see Mann & Bidwell, 1999; Relyea, 2005a, 2005b; Relyea & Jones, 2009). Therefore, we cannot discard other possible harmful effects of herbicides on freshwater communities.

The insecticide fipronil caused a massive reduction in the abundance of aquatic and semiaquatic insects in ‘sugarcane’ ponds, as we expected (H3.1). The fipronil concentration necessary to kill half of a population of chironomid midges in eight days (*i.e.* EC50) is about 3.7 ug L^−1^ (Pinto et al., 2021), a concentration more than four times lower than the one found in ‘sugarcane’ ponds after the fipronil pulse. Amphibians, however, have an estimated lethal concentration (*i.e.* LC50) higher than 800mg L^−1^ (ELG Espíndola, *pers. com*), a concentration more than 10 thousand times higher than our manipulated concentration. Indeed, amphibian abundance was not affected right after the fipronil application. In fact, as we expected, we observed an increase in amphibian abundance one month after the pesticide pulse (*i.e.*, 84-days survey), likely because of the reduced predation pressure from predatory insects. Similar effects have been observed in other experimental work conducted in more controlled mesocosm experiments with different insecticides (*i.e.*, carbaryl: Relyea, 2005, malathion: Relyea et al., 2005, endosulfan: Rohr & Crumrine, 2005). Therefore, we believe these results can be generalized to other monocultures. More importantly, they might even be stronger in crops that, different from sugarcane, heavily relies on the use of insecticides. Soybean fields, for example, are typically subject to three to four applications of insecticides, for a combination of 13 active ingredients within a crop cycle that typically lasts less than four months (Schiesari et al., 2013; Schiesari & Grillitsch, 2011). Indeed, similar patterns of high mortality of predatory insects and a higher abundance of amphibians have been observed in soybean fields, when compared to forest or savanna habitats (Negri, 2015). This is not to say that insecticide applications in agricultural fields may not be detrimental to anuran larvae in any way. For instance, (Relyea & Diecks, 2008) found that insecticide malathion can indirectly reduce the time to metamorphose in amphibians, making them more vulnerable to pond drying. Rather, we argue that the indirect positive effect of insecticides through reduced predation might have a greater short-term consequence on amphibian abundances than possible negative direct or indirect effects.

We also expected that the community recovery after pesticide application, that is, recolonization by aquatic insects and reduction in amphibian abundances, would happen faster in less isolated communities (H3.2), which we did not observe. We observed the same effects for the ‘sugarcane’ treatment in all three isolation distances. We believe this might have happened because the most important amphibian predators, dragonflies (Heyer et al., 1975; Wellborn et al., 1996), which are known to be good dispersers (McCauley, 2006; McCauley et al., 2008), may have been equally able to recolonize ‘sugarcane’ ponds regardless of the isolation treatment. Therefore, the consequences of predator recolonization to amphibian abundances might have been similar in all isolation distances.

By the end of the rainy season, the effects of all our treatments were weakened or even reversed. At this stage of the community assembly, most predatory and non-predatory insects exhibited negative responses to isolation, even though this effect was not strong enough to affect their total abundance patterns; sugarcane communities were still exhibiting different community structure patterns than those of ‘savanna’ ponds with some aquatic insects being still negatively affected; and pasture communities were not significantly different from ‘savanna’ or ‘sugarcane’ ones anymore when compared to the previous survey. This last pattern is surprising considering that we kept adding fertilizers to both pasture and ‘sugarcane’ ponds up to the end of the experiment, thus, one would expect that any differences caused by nutrient addition would have been stronger at the end of the experiment. However, predatory insects, which were the only taxa that seemed to have responded to nutrient additions, were generally less abundant by the end of the rainy season, if compared to our previous surveys. Amphibians followed a similar pattern. Therefore, even though agrochemical application only had its strongest effects in the middle of the rainy season, they are absolutely relevant in temporary ponds given that their very existence is synchronized with pesticide and fertilizer application in sugar cane fields and other rainfed crops, such as soybean and maize. Additionally, some of the most abundant species observed in our mesocosms have aquatic life cycles of less than three months, such as mosquitoes, midges, dragonflies, and amphibians (Ciota et al., 2014; Hamada et al., 2014; Nebeker, 1973; Oliver, 1971). Thus, the processes happening at the peak (*i.e.*, December and January) of the rainy season can strongly affect patterns of adult insect and amphibian emergence, and therefore, regional abundance patterns. These patterns together highlight the importance of transient states of community structure (see Fukami & Nakajima, 2011) in temporary ponds, especially those observed at the peak of the rainy season (*i.e.* 84-days survey). It is also an indication that seasonal population dynamics likely play an important role in how both agrochemicals and spatial isolation affect community patterns.

## CONCLUSION

Our experiment successfully simulated the assembly and reassembly processes of temporary ponds under a full cycle of agrochemical application similar to what is observed in native habitats, pasture, and sugarcane fields. Thus, we show the net consequences of chemical intensification on freshwater insect and amphibian communities. Specifically, we show that, because insecticide and fertilizer application frequently overlap with the rainy season, it can strongly affect freshwater communities in temporary ponds formed close to or embedded in pastures and crop fields. Nutrient enrichment in ‘pasture’ ponds can potentially favor predatory insects through *bottom-up* effects, whereas the use of insecticides in sugarcane fields may strongly decrease aquatic insect abundances, which is followed by an increase in amphibian abundance. Furthermore, we found no evidence that the effects of chemical intensification are dependent on simple isolation by distance, likely because the most abundant predatory insects and amphibians in our study system were not affected by isolation. However, it is important to acknowledge that different land use types might impose different dispersal limitations to different freshwater taxa (*e.g.* Carlson et al., 2016; Hansen et al., 2019), possibly affecting how communities respond to local selective pressures. We also must emphasize that here we focused only on the net effects of the use of agrochemicals on community structure based on species abundance patterns. While our results are important in understanding the consequences of land use intensification to freshwater biodiversity, a full understanding of this process must also consider other aspects of land use intensification, such as environmental complexity and heterogeneity both between and within ponds in different land use types (*e.g*. Schiesari & Corrêa, 2016). In addition to community patterns, we acknowledge that possible behavioral, physiological, and morphological changes that may be caused by pesticide exposure (*e.g.* Freitas et al., 2019; Pinto et al., 2021) could result in important long-term changes in regional metacommunity patterns, however, such effects fall beyond the scope of this study.

## ACKNOWLEDGMENTS

We thank the EESB staff for assistance in pond construction and Luis Vicente P. Cavalaro, Bianca S. Valente, Débora Negrão, Fernanda Simioni, Thais Issi, Cauê Machado, Gabriel Yoneta Monte, João Paulo Alencar, Juliana Quagliano, Lorena Batista, Suzana Marte, Tais do Amaral, Jessica Akane, Angelica Moreira, Samuel Elias Vasconcelos Menezes, Gabriel Banov Evora, Gabrielle Peres Tedeschi, Paula Maria Rosa, Maria Julia Lagioto Buzzini and Rafaela Martins for assistance in the community sampling surveys. We thank Tadeu Siqueira and Paulo Inácio Prado for conceptual and statistical advice as members of RMP theses committee, Paulo Roberto Guimarães Junior, Paulo De Marco Júnior and Evaldo Luiz Gaeta Espindola for providing comments and criticism in RMP thesis defense, and Mathew Leibold and the members of the Leibold lab for insightful comments on early versions of the manuscript. We also thank Renata Pardini and Daniel Lahr for providing lab and office space. This study was funded by Fundação de Amparo à Pesquisa do Estado de São Paulo (FAPESP, grant #2015/18790-3; LA Martinelli PI, L Schiesari Co-PI). RM was supported by Ph.D. fellowships from FAPESP (grants #2017/04122-4 and #2018/07714-2) and Coordenação de Aperfeiçoamento de Pessoal de Nível Superior (CAPES). This research followed a design approved by the Ethics Committee of the Escola de Artes, Ciências e Humanidades of the Universidade de São Paulo (CEUA 003/2016) and was conducted in Estação Ecológica de Santa Bárbara under permits of Instituto Florestal (COTEC 553/2017) and ICMBio (ICMBio 17559-7).

