## supplement for "Impacts of agrochemical intensification on the assembly and reassembly of a mainland-island model metacommunity"

### Index

**SUPPLEMENT 1**

*Sugarcane crop cycle and chosen active ingredients*

Sugarcane is planted in the rainy season from October to March. Between 12 and 18 months later, usually in September or October, the first harvest takes place (*i.e.* ´primary cane´ or ´plant cane´), followed by four to five annual harvests by ratoon cropping (´ratoon cane´) after which the field is reformed. Sugarcane planting involves laying stalk stumps along furrows together with NPK fertilizers and frequently an insecticide to control termites and beetles. Other insecticides are applied as needed but are not as prevalent as biological control with parasitoid wasps and the fungus *Metarhizium* is common and effective in sugarcane plantations. Herbicides are sprayed shortly after planting and before sugarcane sprouts (pre-emergence herbicides) and/or a few weeks later (post-emergence herbicides). In subsequent years NPK fertilizers are applied directly on top of the soil, herbicides are either applied over the row or in localized weeds, and insecticides are applied in a shallow line cut over the sugarcane row (Parra *et al.* 2014, Cantarella & Rossetto 2014, R. Rossetto *pers.com.*).

Selected pesticide active ingredients were among the top-selling insecticides and herbicides in the State of São Paulo, and that are registered for use in sugarcane plantations in Brazil (AGROFIT 2020). These were fipronil, a broad-spectrum phenylpyrazole insecticide commonly used for controlling moths, ants, beetles and termites, and 2,4-D, an alkylchlorophenoxy selective systemic herbicide used for controlling broadleaf weeds. Fipronil is the top-selling insecticide registered for use in sugarcane in the State of São Paulo and is also registered for use in Belgium, the Netherlands, USA, and Australia (PPDB 2020). 2,4-D is the second top-selling herbicide registered for use in sugarcane in the state of São Paulo and is also used in Australia and 27 countries of the European Community (PPDB 2020). The toxicity of fipronil is moderate to algae and moderate to high to zooplankton and sediment-dwelling aquatic insects whereas the toxicity of 2,4-D is low to algae, moderate to macrophytes, and low to zooplankton (PPDB 2020).

#### SUPPLEMENT 2

**Table S2.1.** Measured surface pesticide concentration in sugarcane ponds. Both fipronil and 2,4-D were measured by liquid chromatography coupled with tandem mass spectrometry (LC-MS/MS). Application dates were 30-Oct-2017 and 04-Dec-2017 for fipronil and 2,4-D, respectively.

| Date | Concentration (ug $L^{-1}$) | |
|---|---|---|
| | **Fipronil** | **2,4-D** |
| 30-Oct-2017 | 15.6 | |
| 04-Dec-2017 | 2.9 | 337.8 |
| 04-Mar-2018 | <0.059* | <0.406† |

*concentration below detection limit; †concentration below quantification limit.

**SUPPLEMENT 3**

We took water samples of each pond 73 after the beginning of nutrient addition (*i.e.* 108 days after the beginning of the experiment) to measure total nitrogen (TN) and total phosphorus (TP) concentration in superficial water. Water samples were stored frozen at -20 C until TP and TN concentration was measured. We measured TP and TN through the Valderrama (1981) method.

**Table S3.1.** Type II Wald Chi-square tests for TN and TP concentration at 73 days after the beginning of nutrient addition.

| | **Degrees of freedom** | **Chisq** | **p** | **pairwise comparisons** |
|---|---|---|---|---|
| *TN* | | | | |
| **Land Use** | **2** | **16.393** | **<0.001** | (Control = Pasture) ≠ Sugarcane |
| **Isolation** | **2** | **17.677** | **<0.001** | (30 = 120) ≠ 480 |
| Land Use : Isolation | 4 | 3.648 | 0.456 | |
| *TP* | | | | |
| **Land Use** | **2** | **17.625** | **<0.001** | (Control = Pasture) ≠ Sugarcane |
| Isolation | 2 | 0.039 | 0.981 | |
| Land Use : Isolation | 4 | 4.304 | 0.366 | |

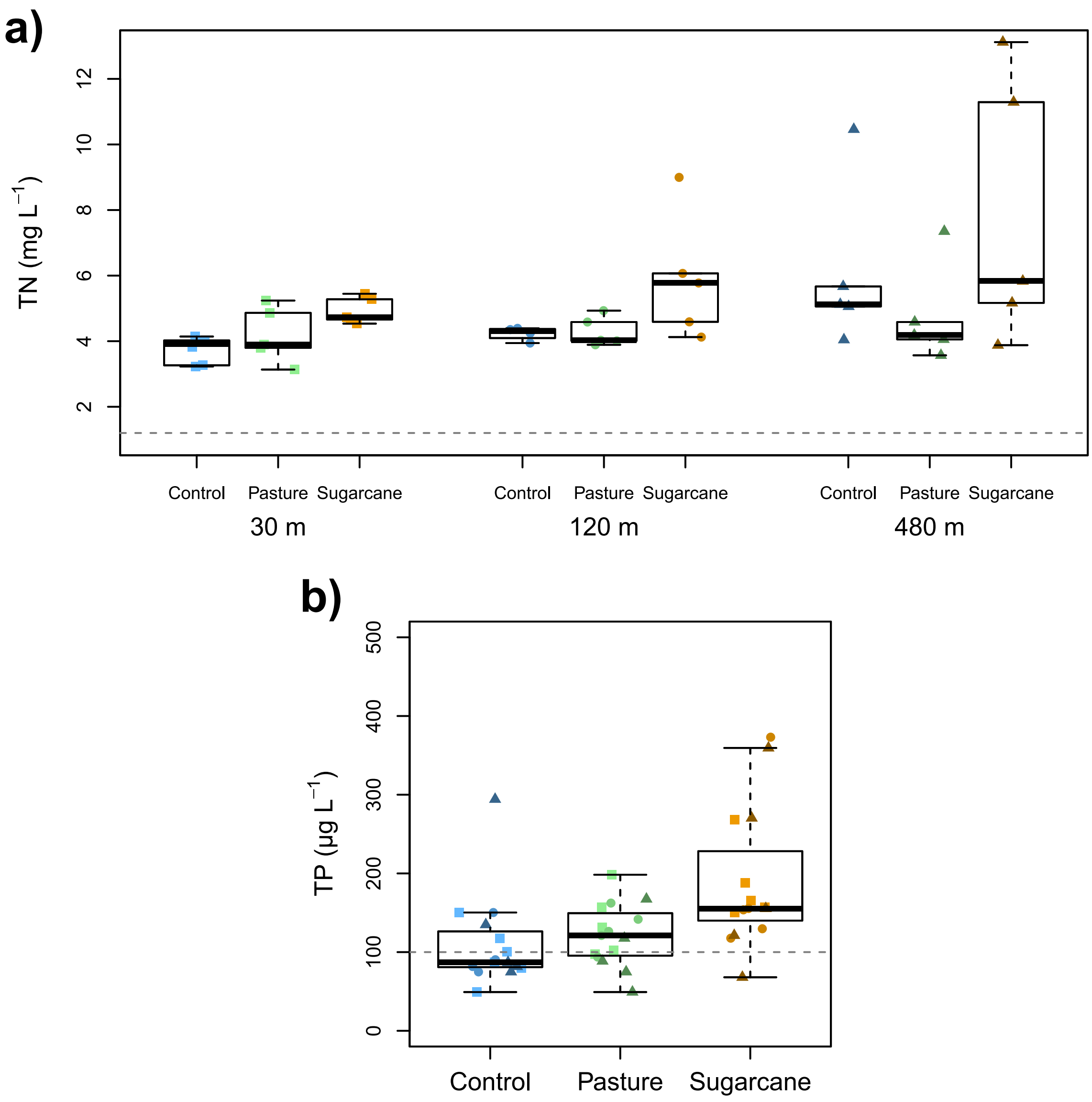

**Figure S3.1.** (a) Total nitrogen and (b) total phosphorus concentration 73 days after the beginning of nutrient additions. Blue, green and yellow symbols are control, pasture and sugarcane treatments, respectively. Squares, circles and triangles are 30m, 120m and 480m of isolation from the source wetland. Grey hatched lines are the thresh hold concentration for hypertrophic lakes (Smith et al., 1999).

**SUPPLEMENT 4**

**Table S4.1.** Summary of deviance analysis for the effect of isolation on total abundance of predatory insects, non-predatory insects, amphibians (Wald Chi-square tests) and community structure (likelihood ratio test) for the 23 days survey. Significant terms are highlighted in bold ($p < 0.05$).

| | Degrees of freedom | Wald Chisq | Deviance | p |
|---|---|---|---|---|
| Predatory Insects | 2 | 0.863 | | 0.649 |
| **Non-Predatory Insects** | **2** | **9.465** | | **0.009** |
| **Amphibians** | **2** | **7.591** | | **0.022** |
| **Community Structure** | **2** | | **249.500** | **0.001** |

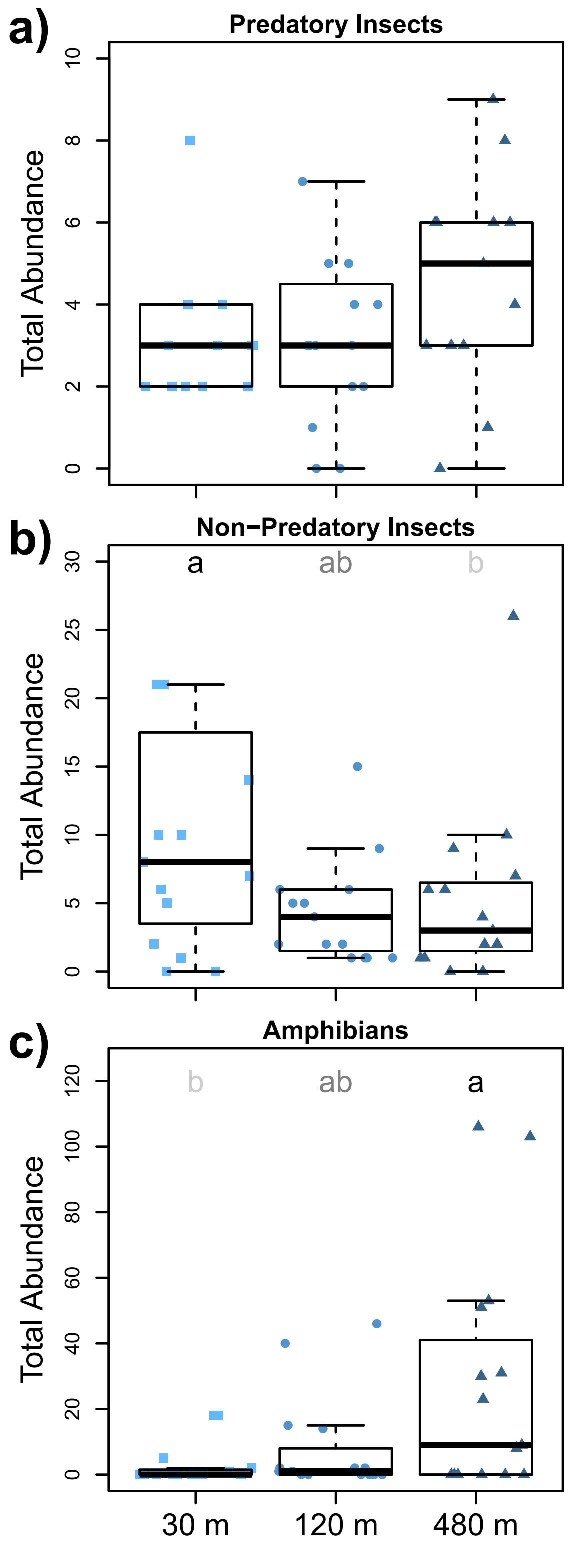

**Figure. S4.1.** Total abundance of predatory insects (a), non-predatory insects (b), and amphibians (c) across the isolation gradient on the 23 days survey. Squares, circles, and triangles are 30 m, 120 m, and 480 m isolation treatments. Different letters represent statistical differences among treatments.

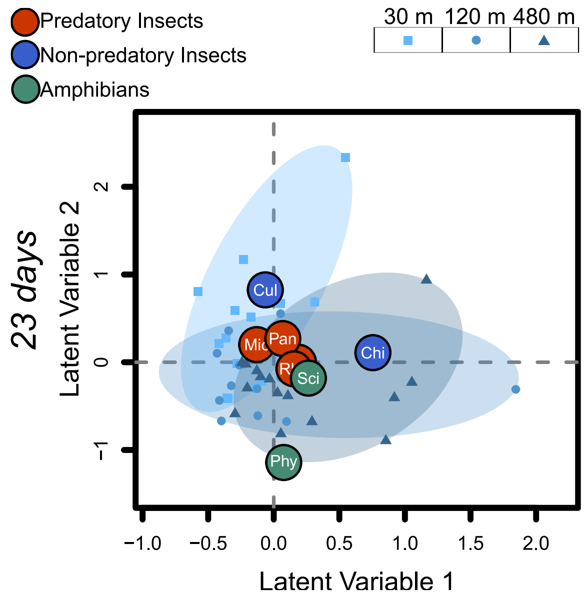

**Figure S4.2.** Model-based unconstrained ordinations showing pond communities (symbols) and species (bubbles) in each of the community surveys (a to c). Abbreviations of names of taxa are provided in supplement 5.

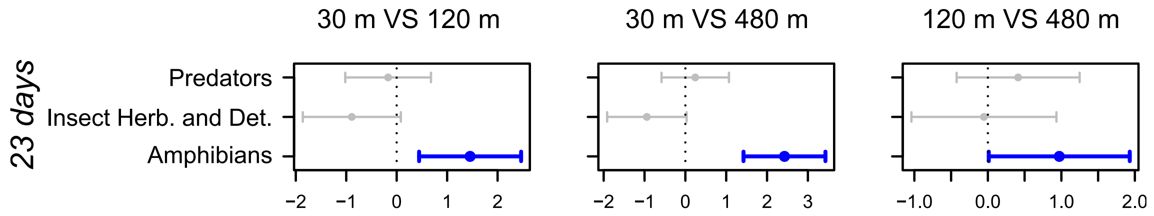

**Figure S4.3.** Pairwise maximum likelihood estimates (dots) and their 95% Confidence intervals (bars) of the differences between isolation (a) and land use (b) treatments for each functional group. Confidence intervals not crossing the zero hatched line were considered significant effects and colored; blue bars represent an increase and red bars a decrease in abundance from the reference treatment (reference VS treatment). These are the mean effects of treatments on different groups of taxa (*i.e.* predatory insects, non-predatory insects, and amphibians). Similar estimates for individual taxa are presented in supplement 7.

**SUPPLEMENT 5**

**Table S2.1.** Family, Order, Class, Trophic level and Total Abundance of all taxa collected during all sampling surveys.

| **Taxa** | **Family** | **Order** | **Class** | **Abbreviation** | **Functional Group** | **Total abundance** | **Reference for Trophic Level** |
|---|---|---|---|---|---|---|---|
| *Scinax* sp. | Hylidae | Anura | Amphibia | Sci | Amphibian | 3079 | Annibale et al. 2019 |
| *Physalaemus nattereri* | Leptodactylidae | Anura | Amphibia | Phy | Amphibian | 1831 | Annibale et al. 2019 |
| *Elachistocleis* sp. | Microhylidae | Anura | Amphibia | Ela | Amphibian | 1 | Annibale et al. 2019 |
| *Rhantus* | Dytiscidae | Coleoptera | Hexapoda | Rha | predator | 46 | Ramírez and Gutiérrez-Fonseca 2014 |
| *Copelatus* | Dytiscidae | Coleoptera | Hexapoda | Cop | predator | 9 | Ramírez & Gutiérrez-Fonseca 2014 |
| *Derovatellus* | Dytiscidae | Coleoptera | Hexapoda | Der | predator | 5 | Ramírez & Gutiérrez-Fonseca 2014 |
| *Bidessonotus* | Dytiscidae | Coleoptera | Hexapoda | Bid | predator | 2 | Ramírez & Gutiérrez-Fonseca 2014 |
| *Laccophilus* | Dytiscidae | Coleoptera | Hexapoda | Lac | predator | 2 | Ramírez & Gutiérrez-Fonseca 2014 |
| *Hypodessus* | Dytiscidae | Coleoptera | Hexapoda | Hyp | predator | 1 | Ramírez & Gutiérrez-Fonseca 2014 |
| *Liodessus* | Dytiscidae | Coleoptera | Hexapoda | Lio | predator | 1 | Ramírez & Gutiérrez-Fonseca 2014 |
| *Heterelmis* | Elmidae | Coleoptera | Hexapoda | Het | herbivore/detritivore | 15 | Ramírez & Gutiérrez-Fonseca 2014 |
| *Berosus* | Hydrophilidae | Coleoptera | Hexapoda | Ber | predator | 390 | Ramírez & Gutiérrez-Fonseca 2014 |

| | | | | | | | |
|---|---|---|---|---|---|---|---|
| *Tropisternus* | Hydrophilidae | Coleoptera | Hexapoda | Tro | predator | 4 | Ramírez & Gutiérrez-Fonseca 2014 |
| *Thermonectus* | Hydrophilidae | Coleoptera | Hexapoda | The | predator | 1 | Ramírez & Gutiérrez-Fonseca 2014 |
| *Hydrocanthus* | Noteridae | Coleoptera | Hexapoda | Hyd | predator | 1 | Ramírez & Gutiérrez-Fonseca 2014 |
| Ceratopogonidae | Ceratopogonidae | Diptera | Hexapoda | Cer | herbivore/detritivore | 40 | Aussel and Linley 1994, Ramírez and Gutiérrez-Fonseca 2014 |
| *Chaoborus* | Chaoboridae | Diptera | Hexapoda | Cha | herbivore/detritivore | 235 | ARCIFA 2000, Ramírez and Gutiérrez-Fonseca 2014 |
| Chironominae | Chironomidae | Diptera | Hexapoda | Chi | herbivore/detritivore | 21560 | Ramírez & Gutiérrez-Fonseca 2014 |
| Tanypodinae | Chironomidae | Diptera | Hexapoda | Tan | herbivore/detritivore | 1885 | Henriques-Oliveira et al. 2003, Ramírez and Gutiérrez-Fonseca 2014 |
| *Culex* | Culicidae | Diptera | Hexapoda | Cul | herbivore/detritivore | 2762 | Ramírez & Gutiérrez-Fonseca 2014 |
| *Callibaetis* | Baetidae | Ephemeroptera | Hexapoda | Cal | herbivore/detritivore | 118 | Ramírez & Gutiérrez-Fonseca 2014 |
| *Caenis* | Caenidae | Ephemeroptera | Hexapoda | Cae | herbivore/detritivore | 374 | Ramírez & Gutiérrez-Fonseca 2014 |
| *Sigara* | Corixidae | Hemiptera | Hexapoda | Sig | predator | 6 | Ramírez & Gutiérrez-Fonseca 2014 |
| *Limnocoris* | Naucoridae | Hemiptera | Hexapoda | Lim | predator | 10 | Ramírez & Gutiérrez-Fonseca 2014 |
| *Pelocoris* | Naucoridae | Hemiptera | Hexapoda | Pel | predator | 1 | Ramírez & Gutiérrez-Fonseca 2014 |
| *Curicta* | Nepidae | Hemiptera | Hexapoda | Cur | predator | 3 | Ramírez & Gutiérrez-Fonseca 2014 |

| | | | | | | | |
|---|---|---|---|---|---|---|---|
| *Buenoa* | Notonectidae | Hemiptera | Hexapoda | Bue | predator | 249 | Ramírez & Gutiérrez-Fonseca 2014 |
| *Notonecta* | Notonectidae | Hemiptera | Hexapoda | Not | predator | 13 | Ramírez & Gutiérrez-Fonseca 2014 |
| *Microvelia* | Veliidae | Hemiptera | Hexapoda | Mic | predator | 1495 | Ramírez & Gutiérrez-Fonseca 2014 |
| *Anax* | Aeshnidae | Odonata | Hexapoda | Ana | predator | 1 | Ramírez & Gutiérrez-Fonseca 2014 |
| *Oxyagrion* | Coenagrionidae | Odonata | Hexapoda | Oxy | predator | 3 | Ramírez & Gutiérrez-Fonseca 2014 |
| *Progomphus* | Gomphidae | Odonata | Hexapoda | Pro | predator | 3 | Ramírez & Gutiérrez-Fonseca 2014 |
| *Pantala* | Libellulidae | Odonata | Hexapoda | Pan | predator | 2419 | Ramírez & Gutiérrez-Fonseca 2014 |
| *Erythrodiplax* | Libellulidae | Odonata | Hexapoda | Ery | predator | 521 | Ramírez & Gutiérrez-Fonseca 2014 |
| *Orthemis* | Libellulidae | Odonata | Hexapoda | Ort | predator | 112 | Ramírez & Gutiérrez-Fonseca 2014 |
| *Tholymis* | Libellulidae | Odonata | Hexapoda | Tho | predator | 5 | Ramírez & Gutiérrez-Fonseca 2014 |

#### SUPPLEMENT 6

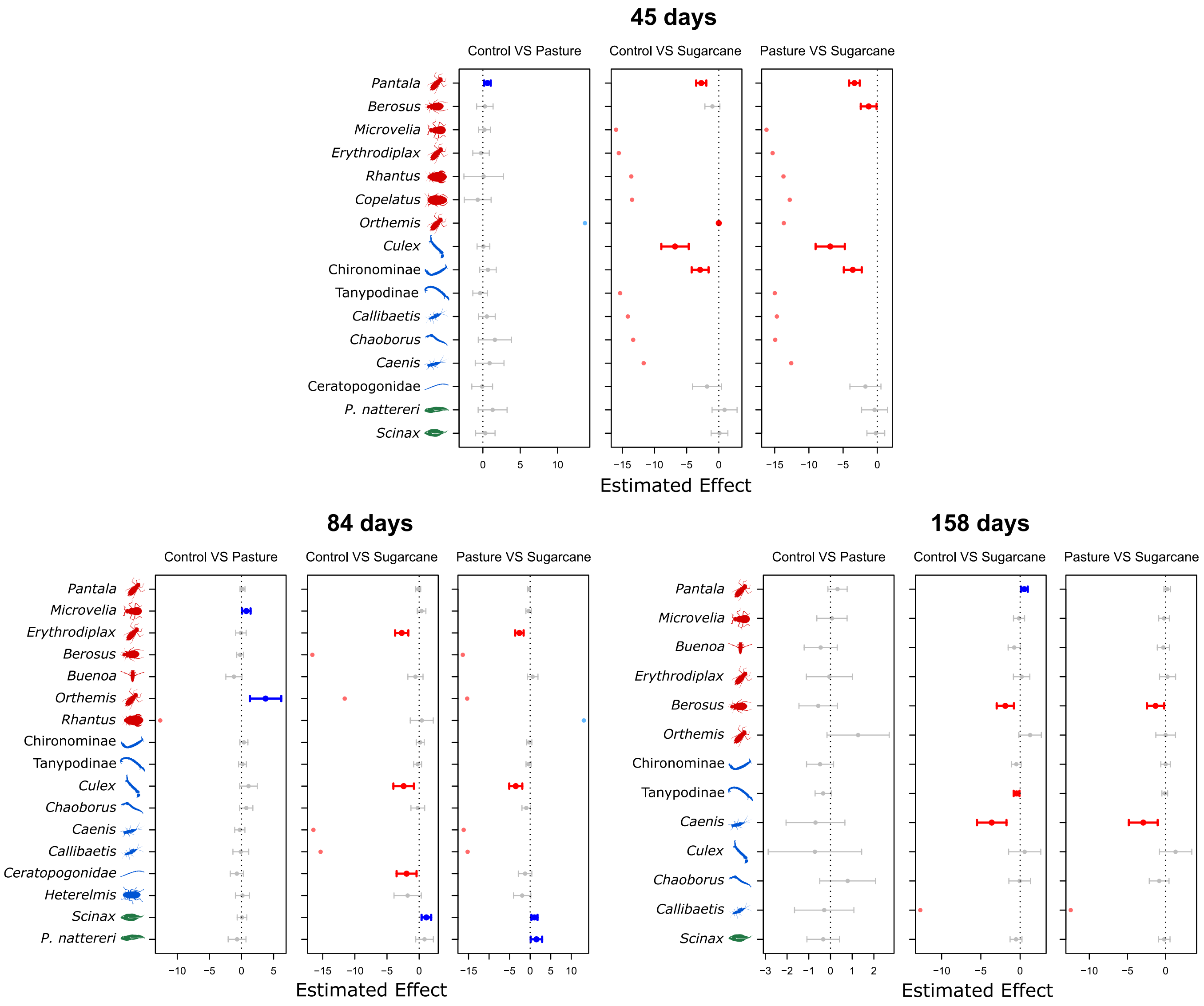

**Figure 6.1.** Confidence intervals for the effect of land use treatments on abundance for each taxon. Taxa are ordered from top to bottom: predatory insects (red cartoons), non-predatory insects (blue cartoons) and amphibians (green cartoons), from most to less abundant. Bars which the 95% confidence interval does not cross the zero-line are colored. Blue bars mean an increase in abundance from the reference treatment. Red bars mean a decrease in abundance from the reference treatment. Lighter blue and red dots mean that a taxon was absent from one of the treatments, thus confidence interval could not be estimated.

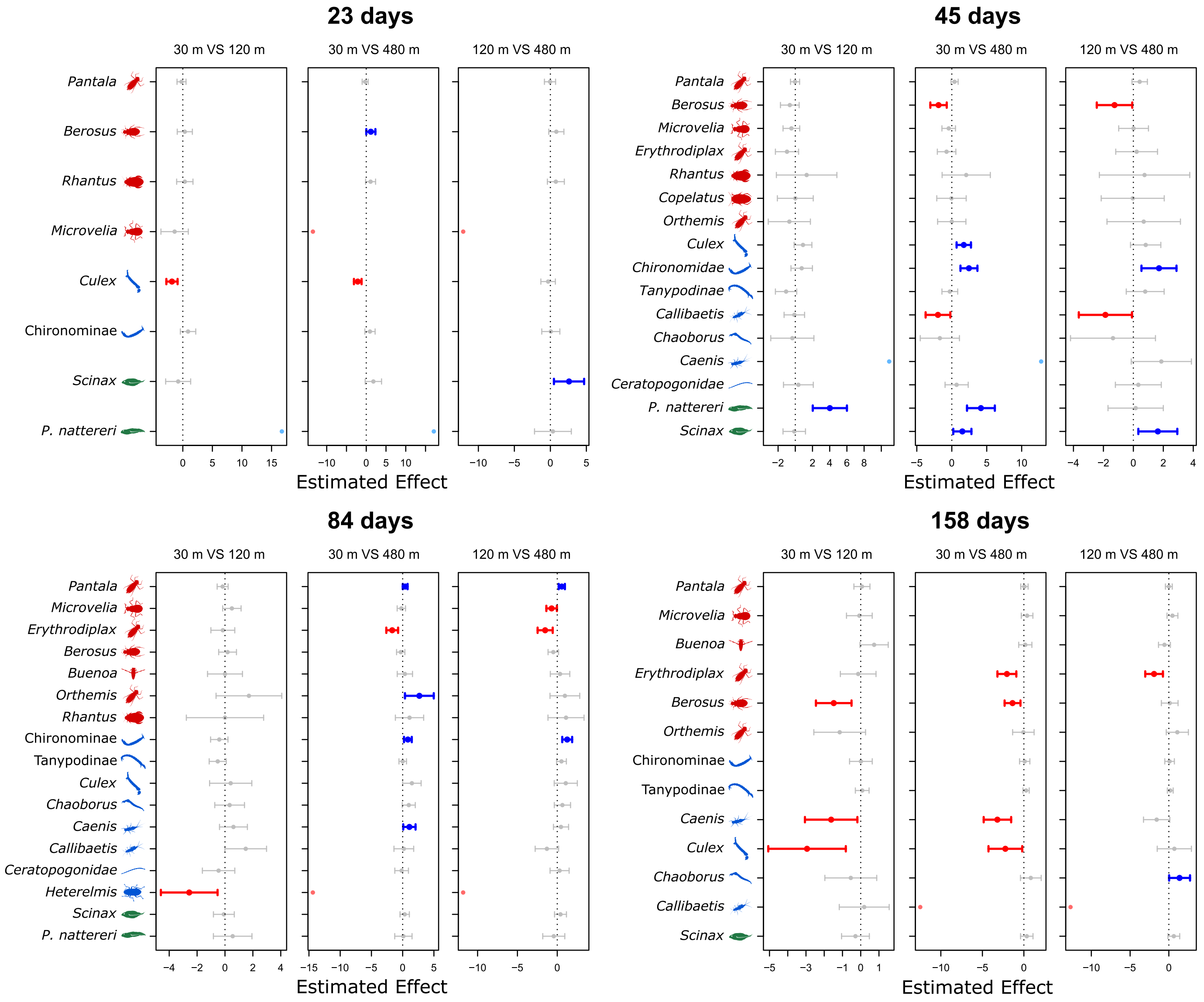

**Figure S6.2.** Confidence intervals for the effect of isolation on abundance for each taxon. Taxa are ordered from top to bottom: predatory insects (red cartoons), non-predatory insects (blue cartoons) and amphibians (green cartoons), from most to less abundant. Bars which the 95% confidence interval does not cross the zero-line are colored. Blue bars mean an increase in abundance from the reference treatment. Red bars mean a decrease in abundance from the reference treatment. Lighter blue and red dots mean that a taxon was absent from one of the treatments, thus confidence interval could not be estimated.
